# Global transcriptome analysis of the *Myxococcus xanthus* multicellular developmental program

**DOI:** 10.1101/564641

**Authors:** J. Muñoz-Dorado, A. Moraleda-Muñoz, F.J. Marcos-Torres, F.J. Contreras-Moreno, A.B. Martin-Cuadrado, J.M. Schrader, P.I. Higgs, J. Pérez

## Abstract

The bacteria *Myxococcus xanthus* exhibit a complex multicellular life cycle. In the presence of nutrients, cells prey cooperatively. Upon starvation, they enter a developmental cycle wherein cells aggregate to produce macroscopic fruiting bodies filled with resistant myxospores. We used RNA-Seq technology to examine the global transcriptome of the 96 h developmental program. This data revealed that many genes were sequentially expressed in discrete modules, with expression peaking during aggregation, in the transition from aggregation to sporulation, or during sporulation. Analysis of genes expressed at each specific time point provided a global framework integrating regulatory factors coordinating motility and differentiation in the developmental program. These data provided insights as to how starving cells obtain energy and precursors necessary for assembly of fruiting bodies and into developmental production of secondary metabolites. This study offers the first global view of developmental transcriptional profiles and provides an important scaffold for future studies.

**IMPACT STATEMENT:** Investigation of global gene expression profiles during formation of the *Myxococcus xanthus* specialized biofilm reveals a genetic regulatory network that coordinates cell motility, differentiation, and secondary metabolite production.

## INTRODUCTION

*Myxococcus xanthus* is a soil-dwelling δ-proteobacterium that exhibits a complex multicellular life cycle with two distinct phases: vegetative growth and starvation-induced development (Muñoz-Dorado et al., 2016). When nutrients are available, cells divide by binary fission to produce a community known as swarm. The *M. xanthus* swarm is predatory (although not obligate) and can digest prokaryotic and eukaryotic microorganisms (Pérez et al., 2016). Upon starvation, cells in the swarm enter a developmental program, during which cells migrate into aggregation centers and climb on top of each other to build macroscopic structures termed fruiting bodies. To form fruiting bodies, starving cells glide on solid surfaces by using two mechanistically distinct motility systems, known as A-(adventurous) and S-(social) motility, which allow individual cell movement or group movement that requires cell-cell contact, respectively (Mauriello et al., 2010; Nan et al., 2014; Islam and Mignot, 2015; Chang et al., 2016; Schumacher and Søgaard-Andersen, 2017). After completion of aggregation (24 h post-starvation), cells differentiate into environmentally resistant myxospores, which are embedded in a complex extracellular matrix (Figure 1). Each fruiting body contains ≈10^5^-10^6^ myxospores. Interestingly, only ≈10% of the starving population become myxospores (O’Connor and Zusman, 1991a) as most cells (around 60%) undergo programmed cell death, most likely to provide the rest of the population enough nutrients to successfully build fruiting bodies (Wireman and Dworkin, 1977; Nariya and Inouye, 2008). The remaining cells differentiate into a persister-like state, termed peripheral rods (PR) which surround the fruiting bodies (O’Connor and Zusman, 1991a, b, c). While PRs are morphologically similar to vegetative cells, myxospores are coccoid and are surrounded by a thick coat mainly consisting of polysaccharides (Müller et al., 2010, 2012; Holkenbrink et al., 2014). Myxospores can germinate when nutrients are available, and collective germination of myxospores from a fruiting body generates a small swarm that facilitates cooperative feeding.

**Figure 1.**
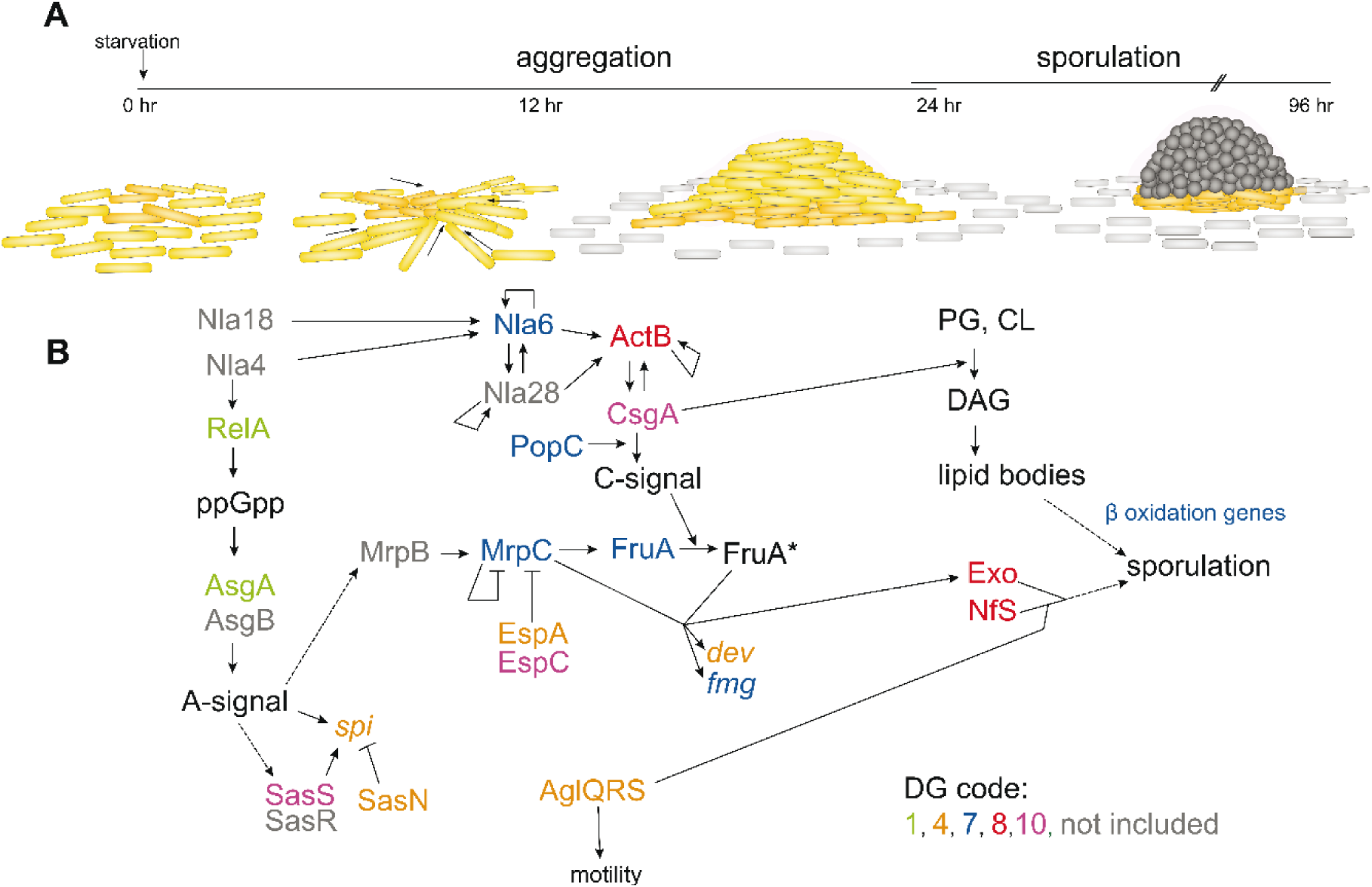
Overview of the *M. xanthus* developmental program and the associated regulatory network. A: Time line of the developmental program indicating aggregation and sporulation phases (top) and schematics of developmental events (bottom). *M. xanthus* cells (yellow rods) aggregate into mounds and then differentiate into resistant spores (grey circles) to produce mature fruiting bodies. Peripheral rods (grey rods) remain outside of the fruiting bodies as a distinct differentiated state. Cells undergoing lysis are not depicted. B: The regulatory pathway for certain well-defined regulatory proteins and signals discussed in the text (see references therein). The developmental group in which the genes were identified is indicated in the legend. Not included: genes not included in the DGs. PG: phosphatidylglycerol; CL: caridiolypin; DAG: diacylglycerol; FruA*: activated FruA.

The developmental program is directed by a sophisticated, but not completely defined, genetic regulatory network which is coupled to a series of intra-and extra-cellular cues. The first cue is starvation, which triggers accumulation of cyclic-di-GMP and, via the stringent response, guanosine penta-and tetraphosphate [(p)ppGpp] inside the cells. These global signals somehow activate four master cascade modules (Nla24, Mrp, FruA, and bacterial enhancer-binding proteins [bEBPs]), which interconnect to control the correct timing of gene expression (Kroos, 2016). Proper progression of development requires intercellular communication, wherein cells produce and transmit five sequential extracellular signals, named A, B, C, D, and E (Bretl and Kirby, 2016).

Although much knowledge has been generated in the last 40 years about the *M. xanthus* developmental cycle, especially with respect to signaling and gene regulatory networks, we are far from having an overall picture of all the events that occur during aggregation and sporulation. Here, we used RNA-Seq technology to measure changes in transcript abundance at 7 time points during *M. xanthus* development. We found that 9.6% of *M. xanthus* genes (1415/7229) had statistically significant changes in transcript abundance during development. Genes in developmentally important pathways were coordinately expressed at the same stage of development. These data and analyses provide, for the first time, a comprehensive view of the transcriptional regulatory patterns which drive the multicellular developmental program of this myxobacterium, offering an essential tool for future investigations.

## RESULTS AND DISCUSSION

### Global transcriptome analysis of the developmental program by RNA-Seq

Global gene expression patterns were examined by RNA-Seq analysis of the wild-type *M. xanthus* strain, DK1622, developed on nutrient limited CF agar plates. RNA was harvested from two independent biological replicates at 0, 6, 12, 24, 48, 72, and 96 h of development, reverse transcribed to cDNA, and sequenced by Illumina methodology (Materials and Methods). On average, 54.72 million read pairs and an average coverage of 591X was obtained. After removing the ribosomal sequences (about 98% of the reads), the genome coverage varied from 5.52 to 14.18X (median of 10.49X), enough reads to provide an adequate coverage of the mRNA fraction. The two sample-replicates showed a high degree of concordance in gene expression (R^2^ correlation >0.98), with the exception of 24-h samples (R^2^ correlation = 0.80). The median of both values was utilized for further analysis (Table 1 and Table 1—Source Data 1).

**Table 1.** Statistical analysis of the *Myxococcus xanthus* DK1622 transcriptome raw data. Data for each of the replicas at 0, 6, 12, 24, 48, 72 and 96 hours of development are shown.

| Sample name | #Gb | #mapped reads | #rRNA-reads | #clean reads (Non-rRNA) | %rRNA rate | coverage (x) | R<sup>2</sup> correlation |
| --- | --- | --- | --- | --- | --- | --- | --- |
| WT_0_1 | 5.70 | 61906718 | 60713758 | 1192960 | 98.07 | 13.05 | 0.99 |
| WT_0_2 | 5.62 | 61117544 | 59826984 | 1290560 | 97.89 | 14.12 |  |
| WT_6_1 | 5.02 | 54468281 | 53756387 | 711894 | 98.69 | 7.79 | 1.00 |
| WT_6_2 | 4.80 | 52436258 | 51759879 | 676379 | 98.71 | 7.40 |  |
| WT_12_1 | 5.04 | 54054132 | 53271826 | 782306 | 98.55 | 8.56 | 0.99 |
| WT_12_2 | 5.37 | 57646891 | 56798096 | 848795 | 98.53 | 9.29 |  |
| WT_24_1 | 3.09 | 33003343 | 32498394 | 504949 | 98.47 | 5.52 | 0.80 |
| WT_24_2 | 5.38 | 53962216 | 53018166 | 944050 | 98.25 | 10.33 |  |
| WT_48_1 | 6.32 | 62796702 | 61500435 | 1296267 | 97.94 | 14.18 | 0.99 |
| WT_48_2 | 5.14 | 34693098 | 34020090 | 673008 | 98.06 | 7.36 |  |
| WT_72_1 | 6.43 | 63625946 | 62530758 | 1095188 | 98.28 | 11.98 | 0.99 |
| WT_72_2 | 5.20 | 51866662 | 50779793 | 1086869 | 97.90 | 11.89 |  |
| WT_96_1 | 6.33 | 63255801 | 62039033 | 1216768 | 98.08 | 13.31 | 0.99 |
| WT_96_2 | 6.11 | 61187496 | 60080963 | 1106533 | 98.19 | 12.11 |  |

As a first data validation step, the expression profiles of many genes that have been previously characterized from β-galactosidase reporter activity (Kroos et al., 1986; Kuspa et al., 1986) or microarray analyses (Bhat et al., 2014) were compared with this data set. While general agreement was observed, strict comparisons were difficult because these assays differed in wild-type strains, developmental conditions, and time points analyzed. Therefore, we analyzed the expression profiles of two different developmental genes using β-galactosidase transcriptional reporters [*spiA*::Tn5-*lacZ* (strain DK4322) and *fmgE*::Tn5-*lacZ* (strain DK4294)] (Kroos et al., 1986) from cells developed under the same conditions used in this study. Comparison of these β-galactosidase activities to the RNA-Seq data indicated the patterns were similar and in agreement with the previously reported results (Supplementary file 1).

### Gene expression profiles organize into 10 developmental groups

To identify developmentally regulated transcripts with similar expression patterns, genes containing measured RPKM (reads per kilobase pair of transcript per million mapped reads) values for all time points were further analyzed. First, all genes with <50 reads, <2-fold changes in RPKM values, and/or high replicate variability between the two replicate datasets (R^2^ correlation <0.7) were removed (Table 1—Source data 2). 1415/7229 (19.6%) genes passed these filters and genes with similar expression patterns were clustered using the kmeans algorithm (Hoon et al., 2004) into 10 developmental groups (DGs) (Figure 2A and Figure 2A— Source data 1). As shown in Figure 2A, analysis of these transcriptional profiles indicated that although some genes (DG1) are down-regulated throughout development, a number of genes are expressed during growth but are also up-regulated at some stage during development (i.e. DGs 2, 5, 6 and 9). Finally, some clusters are exclusive to development, with peak expression either during aggregation (DGs 3 and 4), the transition from aggregation to sporulation (DG7), or sporulation (DGs 8 and 10).

**Figure 2.**
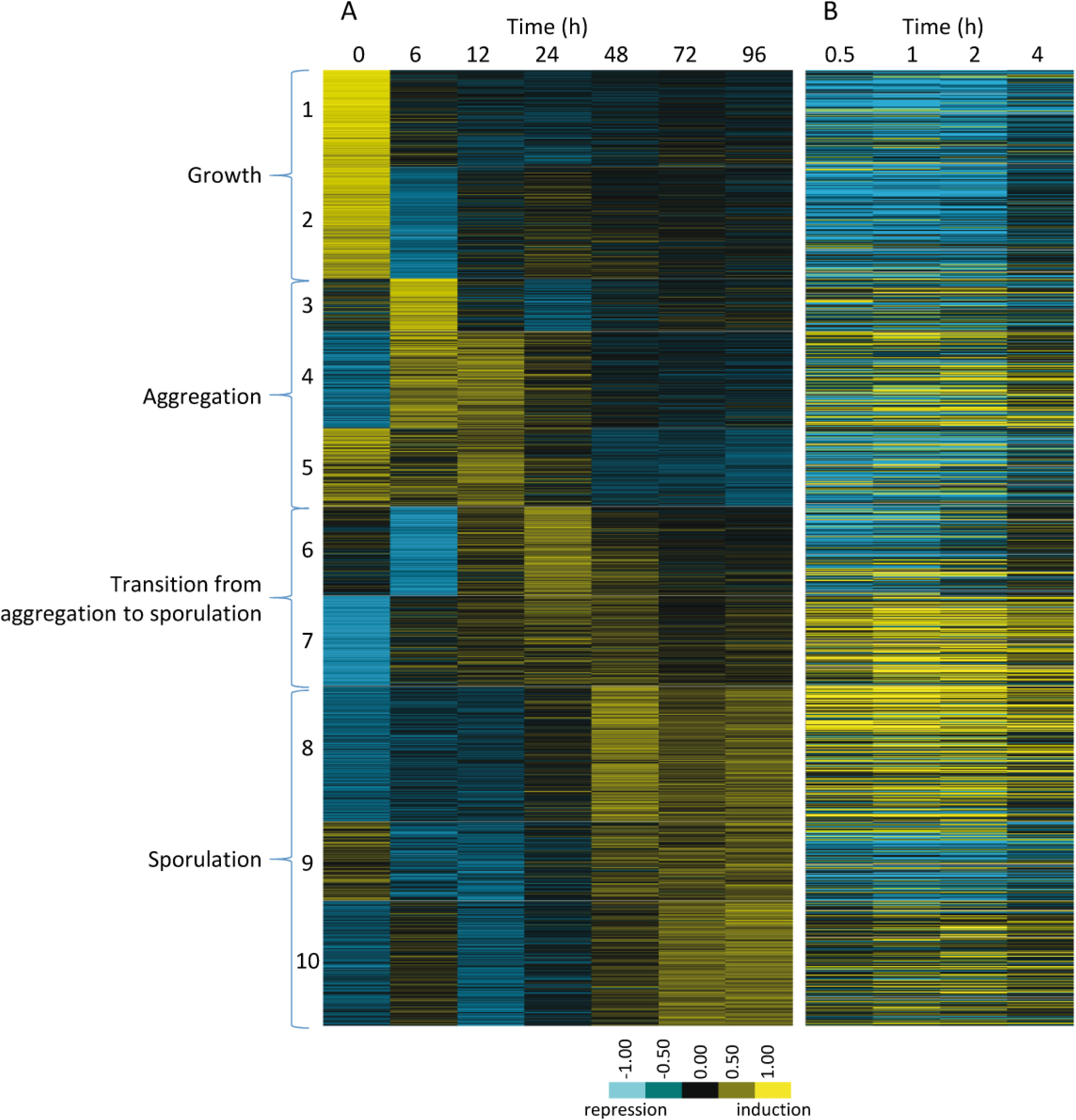
The relative expression profiles of *M. xanthus* genes observed during the developmental program compared to those previously observed during chemical-induction of sporulation. A: Relative expression profiles of significantly regulated genes at the indicated hours after induction of starvation. Genes were clustered into 10 developmental groups based on the time of peak expression and then organized according to the temporal progression of development. Developmental group number and the phase of the developmental program are indicated to the left. B: Relative expression levels of the genes in panel A during the indicated hours after chemical-induction of sporulation (Müller et al., 2010). The position of individual genes in panel B is matched to panel A. Relative expression levels for panels A and B are indicated by color code according to the legend.

### Previously characterized developmental genes can be mapped to each DG

As a first step in analysis and interpretation of the DGs, the expression profile of each DG was analyzed and genes that have been previously described to effect *M. xanthus* development were identified (Figure 2A and Figure 2A—Source data 2). DG1 (144 genes) and DG2 (165 genes) contained genes whose expression decreased at the onset of development with two distinct patterns. Expression of DG1 genes was down-regulated at 6 h and remained at low levels throughout development, while DG2 genes initially dropped drastically but recovered after 6 h, although at lower levels than during growth. Developmentally characterized genes found in these groups include those involved in regulation of the stringent response and induction of the developmental program, such as *relA* and *ndk* (Singer and Kaiser, 1995; Harris et al., 1998; Diodati et al., 2008), *rodK* (Rasmussen et al., 2005), *socE* (Crawford and Shimkets, 2000), and genes responsible for E signal, *esgA* and *esgB* (Downard et al., 1993). However, not all genes thought to regulate entry into development were included in these two DGs. For instance, *nsd*, which governs how cells respond to nutrient availability (Brenner et al., 2004), was in DG4; and *dksA*, which regulates the stringent response (García-Moreno et al., 2009), was in DG5. DG3 included only 78 genes with expression that reached a sharp peak at 6 h and decreased thereafter, such as *pilA*, which codes for the major subunit of the *M. xanthus* pilus filament (Wu and Kaiser, 1995).

DG4, with 143 members, included genes that show low levels of expression during growth, were up-regulated early during development, and then decreased after 24 h. Genes in this group included A signal-related genes *spiA*, MXAN_RS25570, and *sasN* (Cusick, 2015; Kaplan et al., 1991; Kroos et al., 1986; Xu et al., 1998); the *dev* operon (Boysen et al., 2002; Viswanathan et al. 2007); the two-component system *hsfAB* (Gill et al., 1993; Ueki and Inouye, 2002); and genes necessary for production of iso-branched fatty acids (iso-FAs) (Bode et al., 2009; Ring et al., 2006; Bhat et al. 2014). Genes responsible for DK xanthene biosynthesis (Meiser et al., 2006) and many genes involved in A-motility were also located in this group.

DG5 (116 genes) contained genes expressed in growth, but whose expression decreased after 6 h, subsequently increased at 12 h, and then finally decreased again during sporulation, including *fibA* (Kearns et al., 2002; Lee et al., 2011) and many genes involved in A-and S-motility.

DG6 consisted of 132 genes that are expressed at low levels during growth, followed by a decrease in expression at 6 h, and then a dramatic increase at 24 h corresponding to the transition from aggregation to sporulation, such as the gene encoding the protease LonD (Gill et al., 1993; Ueki and Inouye, 2002).

DG7 (134 genes) contained genes that were not expressed during growth, increased from 6 to 12-24 h (aggregation phase), and were maintained at high levels during sporulation. This group contained genes critical for development and related to C-signal and the major developmental regulators *fruA, mrpC*, and *nla6* (Kroos, 2016). Consistently, DG7 also contained genes that are targets of FruA, MrpC, and C signal, such as *fmgA, fmgB, fmgC, fmgD* and *sdeK* (Kroos et al., 1986; Son et al., 2011; Kroos, 2016). Other genes involved in C signaling, such as *popC* (Rolbetzki et al., 2008) and *popD* (Konovalova et al., 2012), were also found in DG7. Finally, this DG also included genes necessary for the β oxidation of lipids, consistent with a proposed role for CsgA (see below) in promoting lipid body production (Bhat et al., 2014; Bullock et al., 2018).

DGs 8, 9, and 10 (with 200, 117, and 186 genes, respectively) contained genes with peak expression levels at the final stages of development, during sporulation. Consistently, many genes up-regulated during chemical induction of sporulation (described below) were found in these DGs (compare Figures 2A and 2B). Additionally, genes involved in secondary metabolite (SM) production and genes encoding type I polyketide synthase (PKS), nonribosomal peptide synthetase (NRPS), and their hybrid (PKS/NRPS) domains were clustered in these DGs. Interestingly, DG10 contained genes that, based on their assigned function, are not expected to be expressed at such a late time. Examples of these genes are *sasS*, which is involved in A signaling (Yang and Kaplan 1997); *csgA*, which encodes the C signal (Shimkets and Rafiee 1990); and *romR*, which is involved in polarity control of motility (Leonardy et al., 2007). However, it is noteworthy that DG10 genes also show high expression at 6 h.

### A-and S-motility genes exhibit different developmental expression profiles

The two phases of the *M. xanthus* lifecycle (predatory growth and developmental aggregation into fruiting bodies) depend on gliding motility (Pérez et al., 2014). Gliding motility is mediated by both A-and S-motility engines and their associated regulatory proteins (Islam and Mignot, 2015; Mercier and Mignot, 2016; Schumacher and Søgaard-Andersen, 2017). Many of the motility genes were developmentally regulated (Figure 3) and most of them were included in DGs 3, 4 and 5 (Figure 2A—Source data 2), indicating that their highest expression was observed during aggregation. Interestingly, the expression profiles of the distinct A-and S-machinery genes clearly differed during development. A-motility genes were up-regulated during early development, while S-motility genes first decreased at 6 h, and then returned to growth levels during aggregation (except for *pilA*). Finally, while most of the genes encoding both motility systems decreased during sporulation (Figure 3), some of them, especially those involved in A-motility, maintained expression during sporulation. These observations are consistent with the repurposing of certain A-motility proteins to function in spore coat assembly machinery (Wartel et al., 2013). Alternatively, as myxospores are non-motile, the high expression levels of A-and S-motility genes that remain after sporulation could be attributed to their expression in the PR subpopulation.

**Figure 3.**
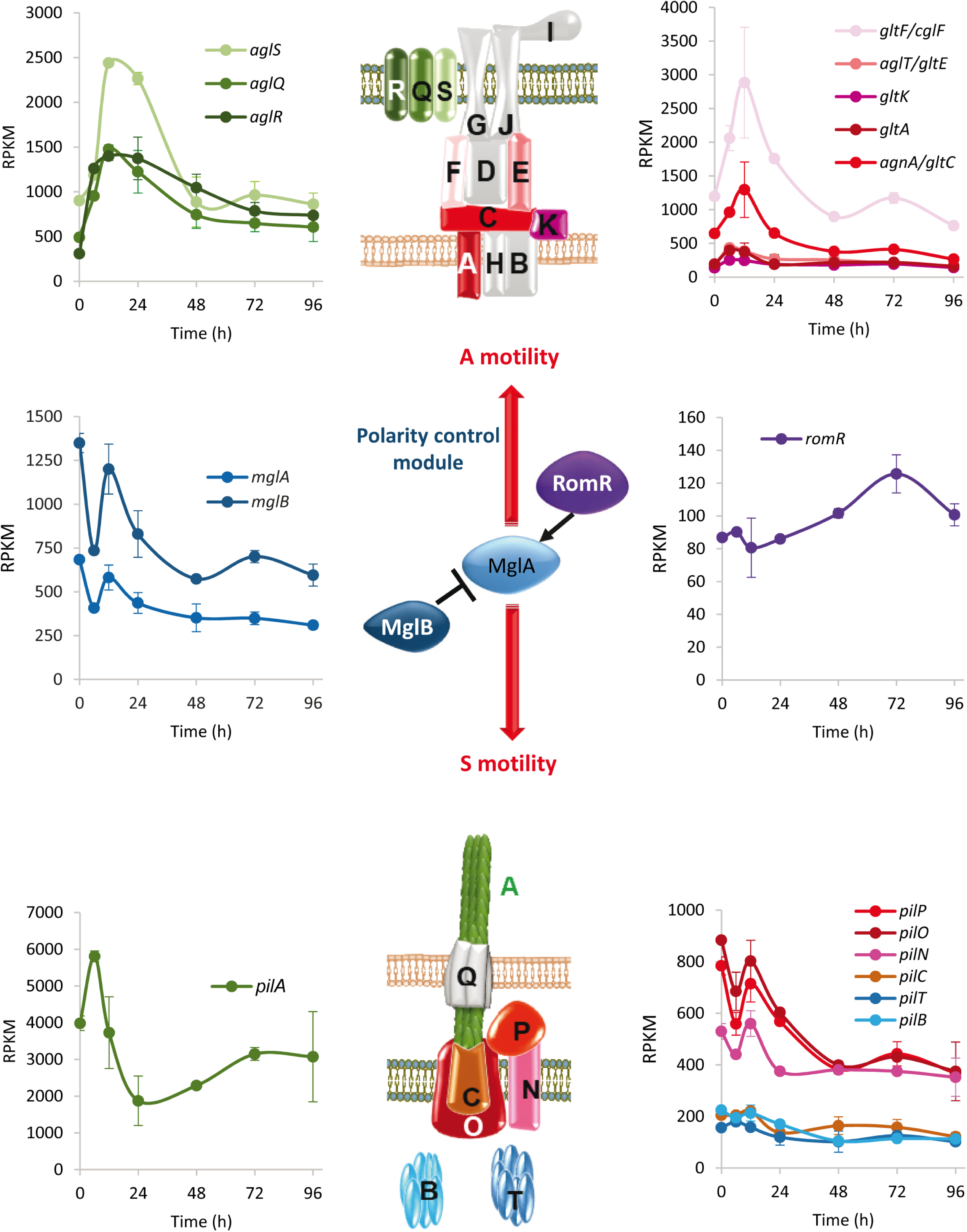
Developmental expression levels of *M. xanthus* motility proteins. Schematic representation of the focal adhesion motor complexes necessary for adventurous (A-) motility (top), the type IV pili motor complexes necessary for social (S-) motility (bottom), and the proteins involved in controlling polarity of both engines (polarity control module; center). The developmental expression levels (RPKM) of significantly regulated motility genes at the indicated times (h) of development. Gene expression profiles are colored to match the proteins depicted in the schematic. Proteins depicted in grey represent genes that were not included in the developmental groups. Error bars indicate standard deviations.

### Gene expression patterns are consistent with use of glycogen and lipid bodies as energy sources

During growth, *M. xanthus* does not appear to consume sugars as carbon or energy sources (Watson and Dworkin, 1968; Bretscher and Kaiser, 1978). Instead, pyruvate, amino acids, and lipids are efficiently utilized which likely directly enter the tricarboxylic acid (TCA) cycle (Bretscher and Kaiser, 1978). It has been long debated as to whether *M. xanthus* utilizes a fully functioning glycolytic pathway (Curtis and Shimkets, 2008). Instead, it was speculated that the pathway may be utilized primarily in the gluconeogenic direction to produce sugar precursors necessary for spore coat production, although homologs of all glycolytic genes and putative sugar transporters genes can be identified in the genome (Youderian et al., 1999; Chavira et al., 2007; Gestin et al., 2013). It is unknown how these pathways contribute to energy production during development when starving cells must synthesize energy currencies (i.e. ATP) over a period of at least three days.

Analysis of the DGs revealed that most genes involved in energy generation (pyruvate dehydrogenase complex, TCA cycle and oxidative phosphorylation) were found in DG2 (Figure 4A). Interestingly, many genes encoding enzymes of the glycolytic/gluconeogenic pathway were up-regulated during development, with most reaching maximum expression levels at the completion of aggregation (24-48 h) (Figure 4B). This up-regulated group included genes encoding homologs of glucokinase (*glkC*; MXAN_RS31480) and phosphofructokinase (*pfkA*; MXAN_RS19530) which are specific for the glycolytic pathway (Figure 4B). These observations are consistent with a transcriptional rewiring of metabolic pathways during the developmental program, perhaps to take advantage of changing carbon/energy sources. For instance, developmental up-regulation of the glycolytic pathway genes may allow developing cells to obtain energy from sugars released from cells undergoing developmental lysis or from glycogen. Glycogen accumulates during late stationary phase/early development, and then disappears prior to sporulation (Nariya and Inouye, 2003). Enzymes predicted to be involved in synthesis of glycogen, such as GlgC (MXAN_RS07405), were found in the DG4 (aggregation phase), while enzymes involved in utilization of glycogen, such as MXAN_RS17870, GlgP (MXAN_RS28270), and MalQ (MXAN_RS31300), appeared to be constitutively expressed (Table 1—Source data 2). The observation that these competing pathways show overlapping expression profiles suggests that regulation of glycogen production/consumption is likely regulated post-transcriptionally, as has been demonstrated by phosphorylation of PfkA by Pkn4 (Nariya and Inouye, 2003).

**Figure 4.**
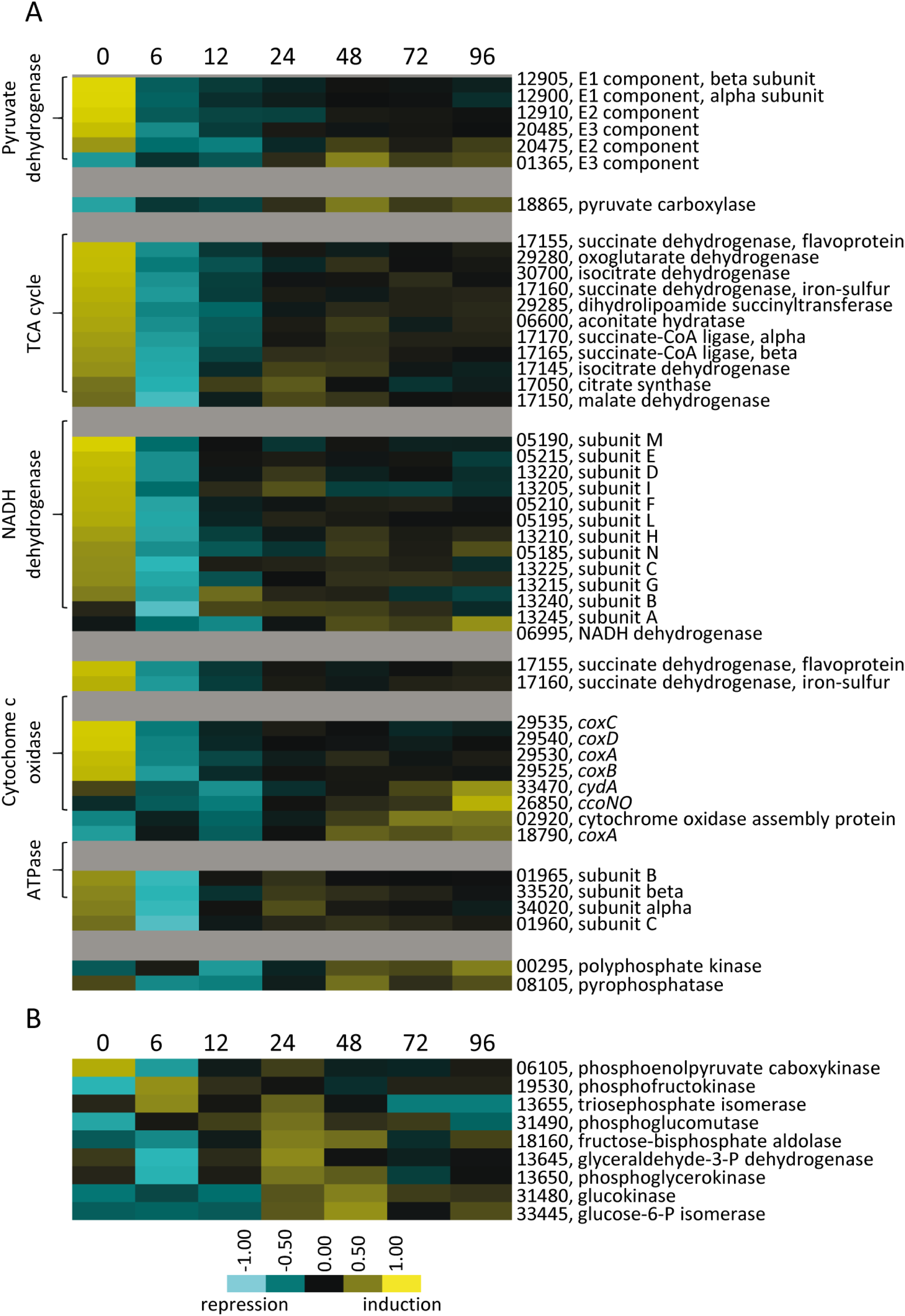
Relative developmental expression profiles of genes involved in energy generation. A: Genes encoding protein homologs for the pyruvate dehydrogenase complex, TCA cycle, and oxidative phosphorylation proteins. B: Genes necessary for glycolysis/gluconeogenesis. Developmental time points in hours are indicated above each panel. Relative expression levels for panels A and B are indicated by color code according to the legend. For simplicity, the MXAN_RS designation was omitted from the locus tag of each gene.

Lipid bodies also accumulate in cells prior to sporulation and it has been proposed that they are later used as an energy source (Hoiczyk et al., 2009). The profiles of genes involved in both straight-and branched-chain primers synthesis and elongation of FAs (Figures 5A and 5B) suggested that some level of lipid synthesis occurs during development. Moreover, genes involved in the alternative pathway to produce isovaleryl-CoA (Bode et al., 2009) were induced (Figures 5A and 5B). With respect to lipid degradation, genes involved in β-oxidation (Bhat et al., 2014) were induced from the onset of development, and reached maximum expression at 24-48 h (Figure 5C). These results and other FA degradation enzyme profiles (Figure 5C) suggest lipids are likely consumed preferably during aggregation and initiation of sporulation, but consumption may be maintained up to 96 h. These data strongly suggest that macromolecules recycled from growth phase or released from lysing cells can be directly used to yield energy, but are also used to synthesize glycogen and lipids that are stored for later consumption.

**Figure 5.**
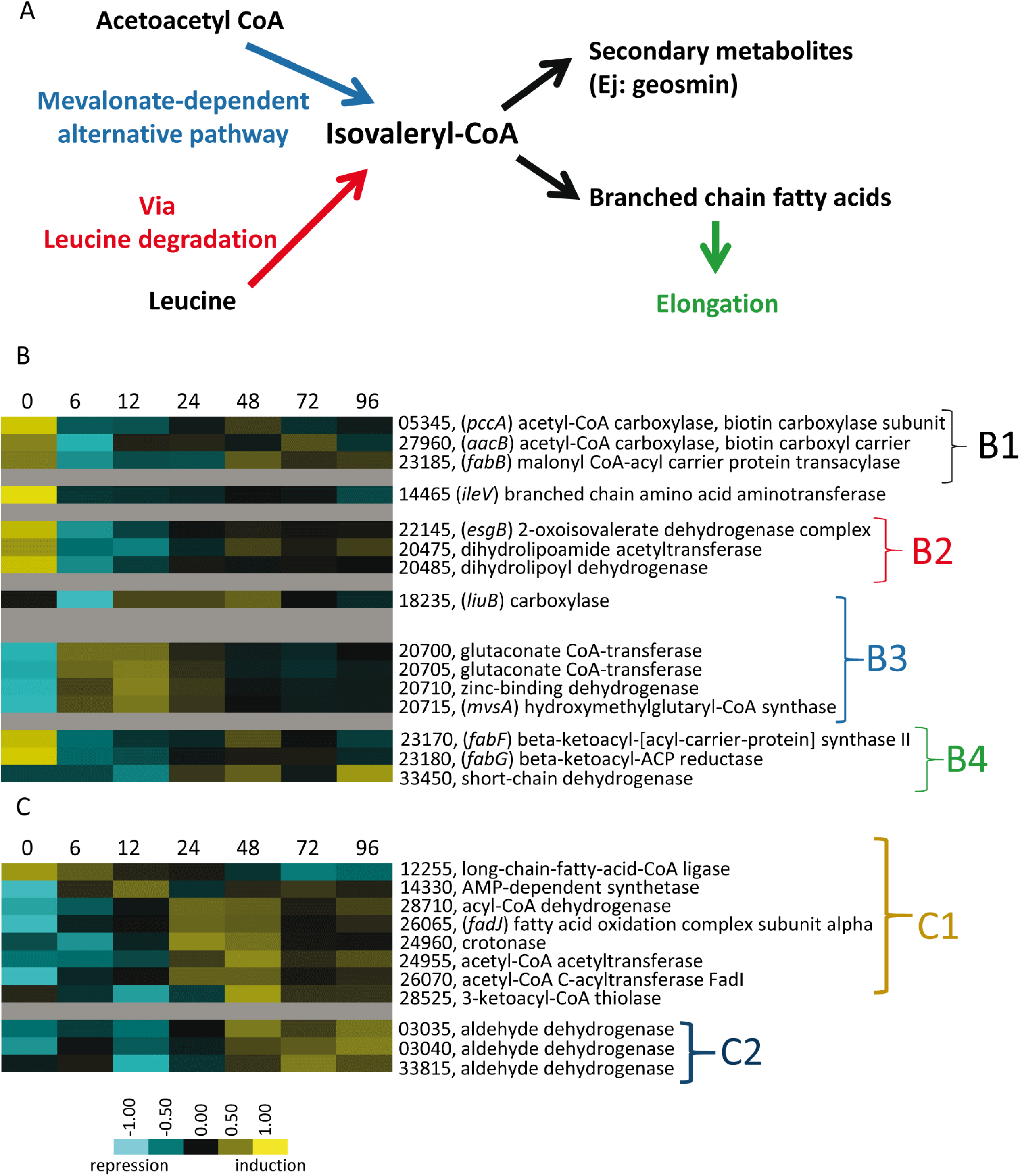
Expression patterns of genes necessary for synthesis and degradation of lipids. A: Simple representation of the *M. xanthus* branched fatty acid metabolic pathways depicting leucine degradation and alternative mevalonate-dependent routes. B: Relative developmental expression profiles of the genes involved in straight-chain and branched-chain fatty acid biosynthesis as designated to the right. Developmental time points in hours are indicated above each panel. B1: Straight-chain fatty acid primer synthesis; B2: Branched-chain fatty acid primer synthesis of isovaleryl-CoA via leucine degradation (*bkd* genes); B3: Branched-chain fatty acids primer synthesis of isovaleryl-CoA via the alternative pathway (mevalonate); B4: Fatty acid elongation. C: Lipid degradation via β oxidation (C1) and other pathways (C2). Relative expression levels for panels B and C are indicated by color code according to the legend. The MXAN_RS designation was omitted from the locus tag of each gene.

### Amino acid and sugar precursors required for developmental macromolecule synthesis are likely released by protein and polysaccharide turnover and gluconeogenesis

In addition to energy, starving cells need a source of sugar precursors to synthesize developmentally specific polysaccharides required for motility (Li et al., 2003), fruiting body encasement (Lux et al., 2005), spore coat synthesis (Kottel et al., 1975; Holkenbrink et al., 2014), and spore resistance (McBride and Zusman, 1989). It has been suggested that these sugars are derived from gluconeogenesis (Youderian et al., 1999). Consistently, our data have revealed that genes encoding enzymes specific for gluconeogenesis, such as phosphoenolpyruvate carboxykinase (MXAN_RS06105) was in DG2, and GlpX (fructose-1,6-bisphosphatase; MXAN_RS21640) was present during growth and throughout development (Table 1—Source data 2). Thus, these observations, as well as those presented above (Figure 4A), indicate that gluconeogenesis contributes to sugar precursor production at various stages during development. Moreover, the observation that four glycosyl hydrolases were specifically up-regulated during development (Figure 6A) suggests the cells may recycle vegetative polysaccharides or scavenge polysaccharides released from cells induced to lyse. These released free monomers could be synthesized into specific developmental polysaccharides by the series of glycosyl transferases that are also developmentally up-regulated (Figure 6B).

**Figure 6.**
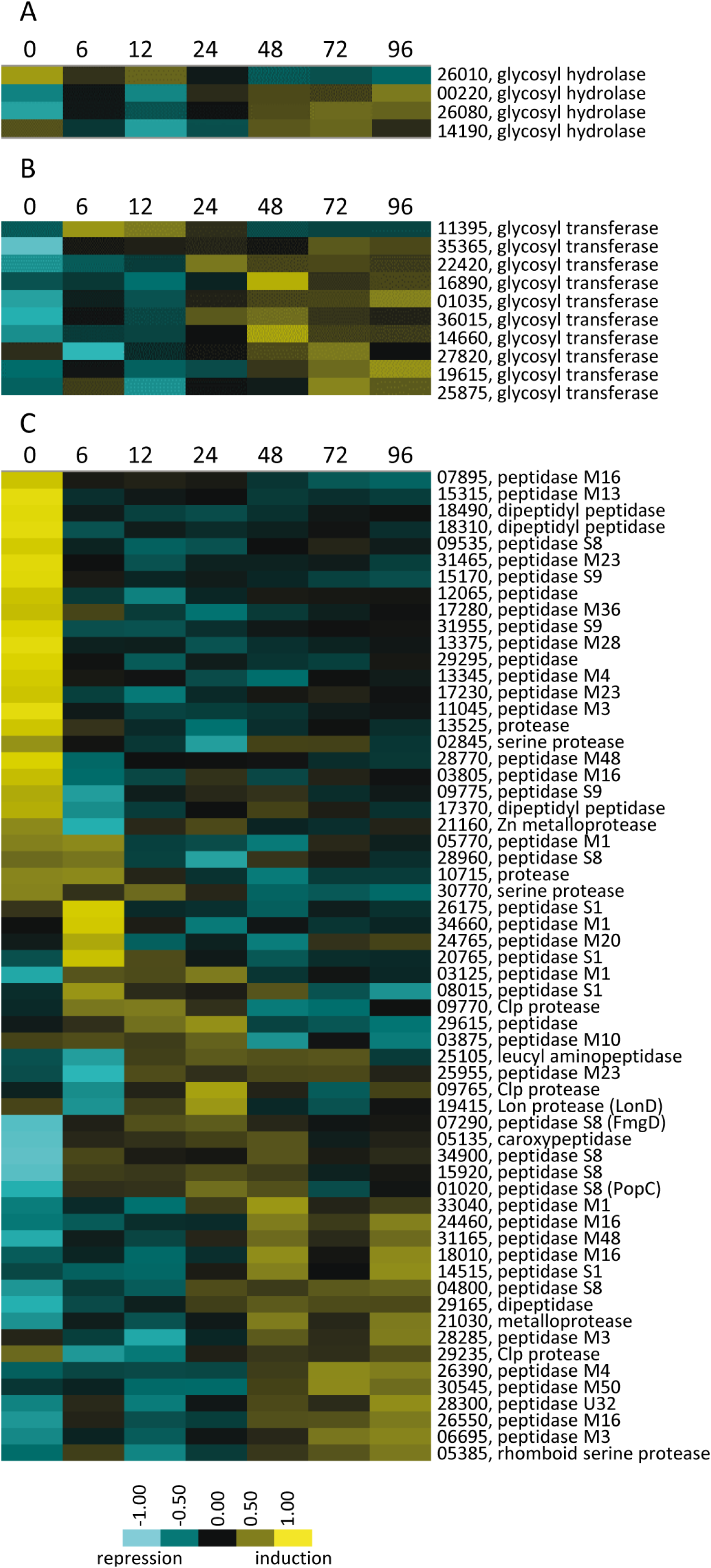
Developmental expression profiles of genes involved in production of polysaccharides and proteins. Relative expression profiles of genes predicted to be necessary for polysaccharide hydrolysis (A), polysaccharide synthesis (B), and encoding proteases and peptidases (C). Developmental time points in hours are indicated above each panel. Relative expression levels for panels A-C are indicated by color code according to the legend at the bottom. The MXAN_RS designation was omitted from the locus tag of each gene.

The developmental program is mainly triggered by amino acid starvation, yet developmentally specific proteins need to be newly synthesized. A source of these amino acids could be released from lysed cells and turnover of proteins that are not required during development. Consistently, 60 proteases appeared in the 10 DGs. 16 of these were specific to growth and down-regulated after 6 h of starvation, while the rest were sequentially up-regulated at different time points during development (Figure 6C). All of these proteases likely release amino acids, but some of them may additionally function in regulatory processes. For example, the protease PopC, which cleaves CsgA p25 to generate the p17 fragment (Rolbetzki et al., 2008) thought to function as the C signal (Lobedanz and Søgaard-Andersen, 2003), is found in DG7.

### Genes involved in secondary metabolism are developmentally up-regulated

Like many of the Myxobacteria, *M. xanthus* produces multiple SMs that facilitate predation during growth (Xiao et al., 2011). The *M. xanthus* genome codes for 18 NRPS, 6 PKS/NRPS, and 22 PKS genes located in regions predicted to be involved in SM synthesis (Korp et al., 2016). Of these, 10 were developmentally up-regulated, and 9/10 were found in DGs 8, 9 and 10. Moreover, an additional 127 genes assigned to the DGs were located in those SM regions (Korp et al., 2016), and 51 of these were also identified in DGs 8, 9, and 10.

Myxovirescine is involved in predation on *Escherichia coli* (Xiao et al., 2011) and, consistently, all the genes involved in its synthesis were expressed during growth. Surprisingly, however, they were also up-regulated during development. Four PKS and one PKS/NRPS myxovirescine synthesis genes (Simunovic et al., 2006) were found in DG10, while the rest of the genes were found in DGs 6, 7, and 8 (Figure 7). Likewise, genes involved in myxoprincomide biosynthesis (Cortina et al., 2012), which is required for effective predation on *Bacillus subtilis* (Müller et al., 2016), as well as genes responsible for geosmin biosynthesis (Bode et al., 2009), were located in DGs 8, 9 and 10 (Figure 2A—Source data 2), suggesting a role in late stages of development. The profiles of genes involved in DK xanthene biosynthesis (Figure 2A—Source data 2), the major yellow pigment of *M. xanthus*, suggested that it may play a role in aggregation in addition to spore maturation (Meiser et al., 2006). Together, these observations suggest that SMs may play previously unknown roles during development. For instance, they may be used to protect developing cells from other microbes in the soil, kill competitors to yield nutrients, or used as signaling molecules. These data and those presented below concerning chemical induced sporulation lead to an intriguing speculation that PRs may produce SMs to defend spores inside the fruiting bodies or to release nutrients from prey to promote germination.

**Figure 7.**
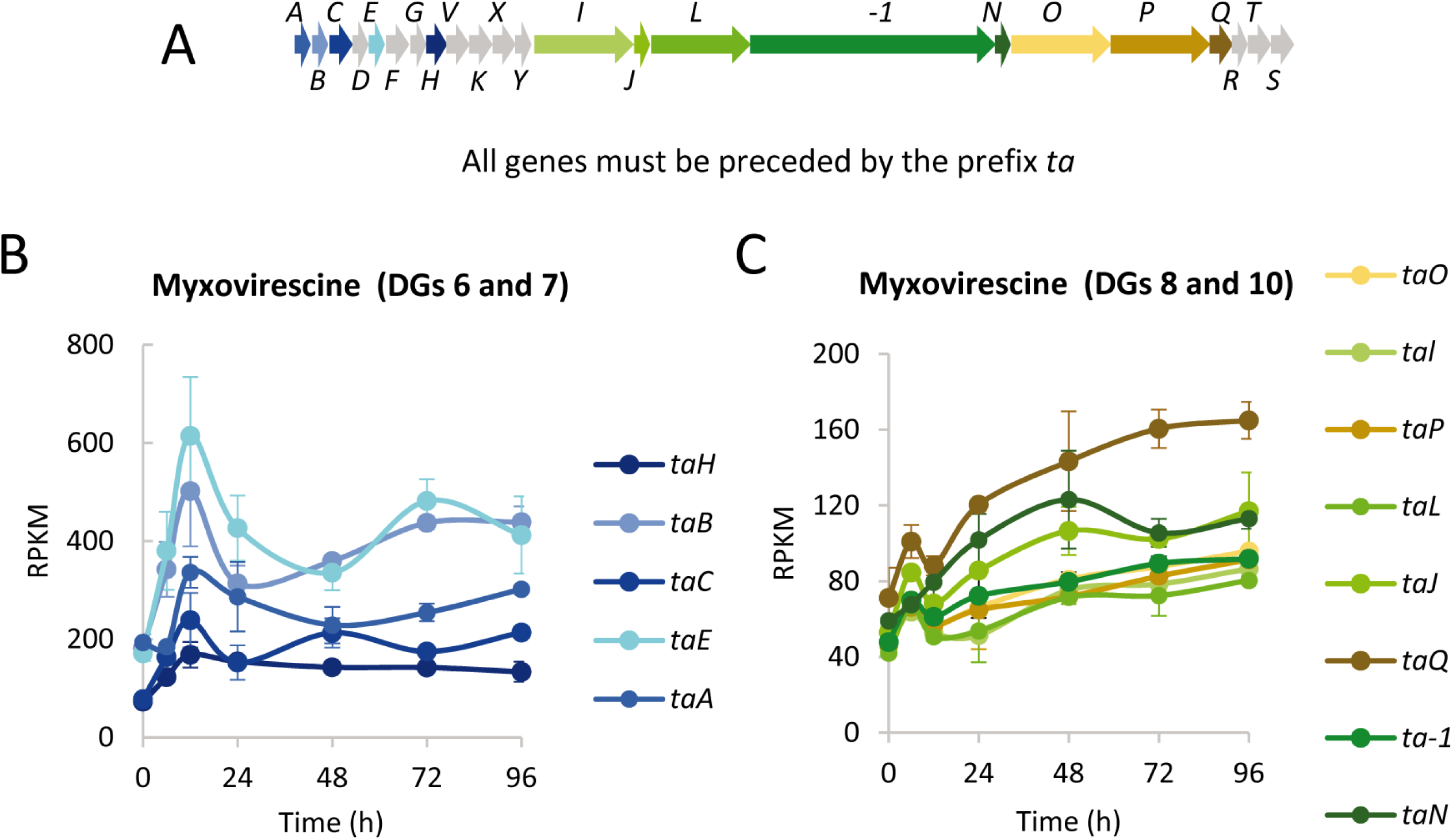
Developmental expression levels of genes involved in myxovirescine (antibiotic TA) biosynthesis. A: Schematic of the *ta* gene cluster. B and C: The developmental expression level (RPKM) of significantly regulated *ta* genes plotted against the indicated developmental time points in hours. Gene expression profiles are colored to match the genes depicted in the schematic. Genes depicted in grey were not included in the developmental groups. Error bars indicate standard deviations.

### Translation may be developmentally rewired

76 genes involved in translation fit the criteria for inclusion in the DGs, which represents 5.4% of the genes included in these 10 DGs. Analysis of genes encoding protein biosynthesis machinery (i.e. ribosomal proteins, initiation, elongation and termination factors, proteins involved in ribosome maturation and modifications, and several aminoacyl-tRNA ligases) revealed that they stop being transcribed at 6 h (Figures 8A and 8B), most likely as the result of the stringent response (Starosta et al., 2014). However, most of them were then up-regulated at later times, reaching a level similar to, or higher than, that observed during growth. A few, including several aminoacyl-tRNA ligases, were mainly expressed during sporulation (Figure 8A). The observation that many ribosomal proteins are differentially regulated during development (Figure 8B) suggests that the ratio among the different ribosomal proteins varies through development, yielding ribosomes with altered protein composition. Consistently, it was previously observed that the protein composition of ribosome complexes purified from growing cells versus myxospores was different (Foster and Parish, 1973). The data presented here confirm these previous results, and expand these observations by providing an overview of how ribosomes may change during progression of development. Another interesting observation was the differences in expression profiles exhibited by duplicated genes of ribosomal proteins. *M. xanthus* encodes two paralogs for proteins S1, S4, S14, L28, and L33 (Table 1—Source data 2). Only one of the two paralogous genes for ribosomal proteins S1 and S14 was found in the DGs for (Figure 2A—Source data 1), suggesting that the ratio between the S1 and S14 paralogs changes during development. Most notably, the S4 paralogs contained similar RPKM during growth, but by 48 h of development the RPKM for MXAN_RS16120 was 10-fold higher than those for MXAN_RS32955 (Figure 8C). Neither of the two paralogs of the 50S subunit were found in the DGs (Table 1—Source data 2).

**Figure 8.**
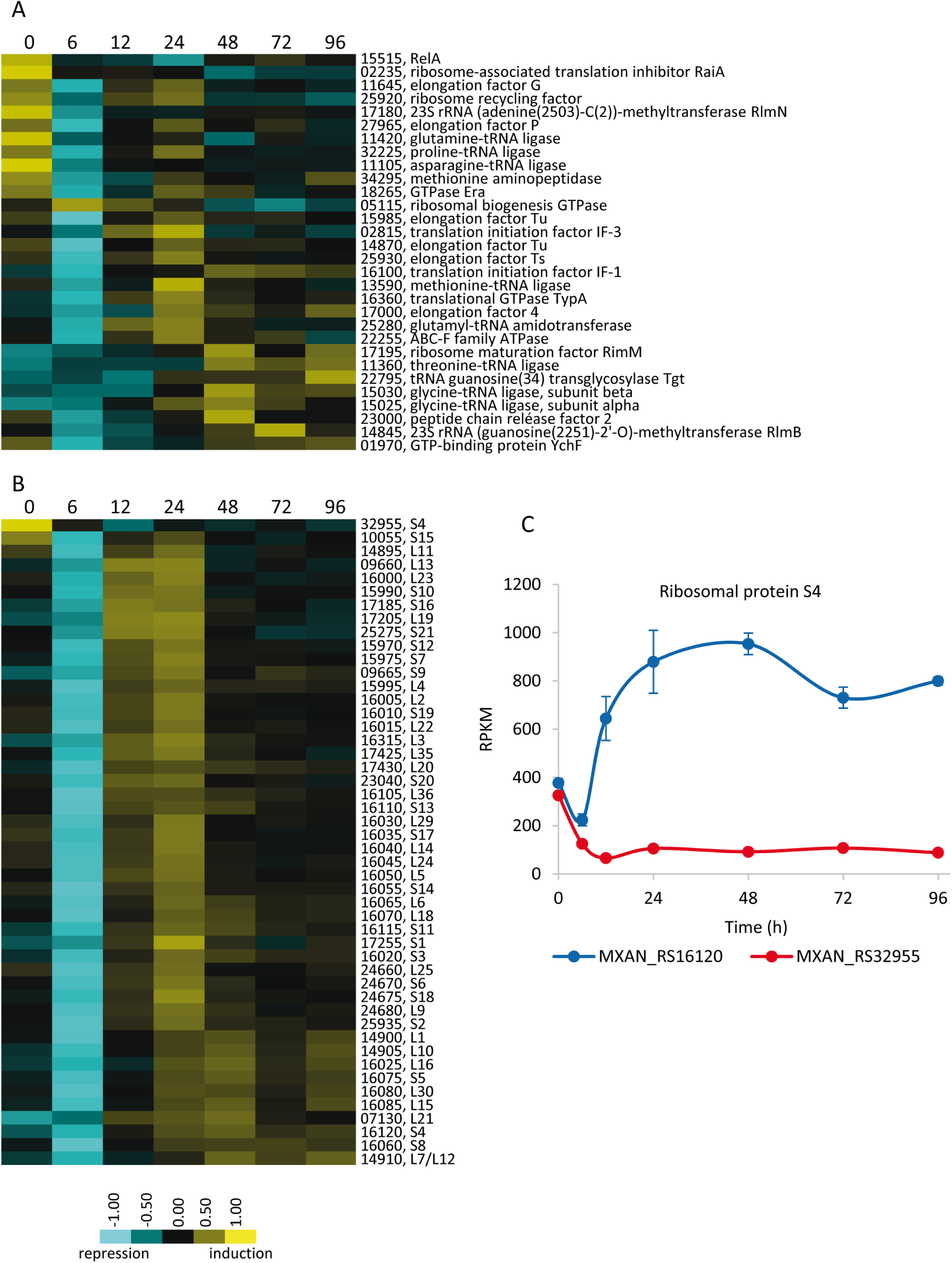
Developmental expression profiles of genes involved in protein production. A: Relative expression levels of genes involved in translation or ribosome assembly. B: Relative expression levels of genes encoding ribosomal proteins. Developmental time points in hours are indicated above each panel. The MXAN_RS designation was omitted from the locus tag of each gene. Relative expression levels for panels A and B are indicated by color code according to the legend at the bottom. C: Developmental expression levels (RPKM) of the paralogous genes encoding protein S4 plotted against developmental time points in hours. Error bars indicate standard deviations.

Some bacteria build alternative ribosomes to improve fitness on different growth conditions by altering the core ribosomal protein stoichiometry and differential expression of paralogous ribosomal proteins (Foster and Parish, 1973; Nanamiya et al., 2004; Hensley et al., 2012; Prisic et al., 2015; Slavov et al., 2015). In addition, some ribosomal proteins also play extraribosomal functions (Warner and McIntosh, 2009). Thus, our data suggest that regulation of translational machinery may play an important role in directing the developmental program. However, we cannot rule out that some of the transcriptional changes observed in the translational machinery could be related to ribosomal hibernation in myxospores and/or PRs (Yoshida and Wada, 2014; Harms et al., 2016; Gohara and Yap, 2018).

### A large interconnected regulatory network controls development

*M. xanthus* encodes a large repertoire of signaling/regulatory proteins presumed necessary to direct and coordinate its multicellular lifecycle in response to extra-and intra-cellular cues. Examples include one-component systems (OCS; i.e. transcriptional regulators that contain a sensing domain), two-component signaling genes (TCS; i.e. sensor histidine kinase [HK] and response regulator [RR] proteins connected by phosphorelay), and serine/threonine protein kinases (STPK). Many of these proteins have been characterized in this bacterium, and in some cases, individual signal-transduction pathways and their exact role in controlling development have been defined (Inouye et al., 2008; Schramm et al., 2012; Muñoz-Dorado et al., 2014; Rajagopalan et al., 2014). However, the data presented here provide for the first time a view of the entire developmental cycle as an integrated system. We observed developmental up-or down-regulation of a larger number of regulatory genes than previously reported; a significant number of these genes showed expression patterns that suggested they likely play important roles in modulation of aggregation and/or sporulation.

21 OCS genes were observed as significantly developmentally regulated, 9 of which were down-regulated. The remaining were differentially up-regulated during development (Figure 9A), including the repressor LexA (Campoy et al., 2003), and the regulators SasN (Xu et al., 1998) and MrpC (Sun and Shi, 2001; Ueki and Inouye, 2003; McLaughlin et al., 2018).

**Figure 9.**
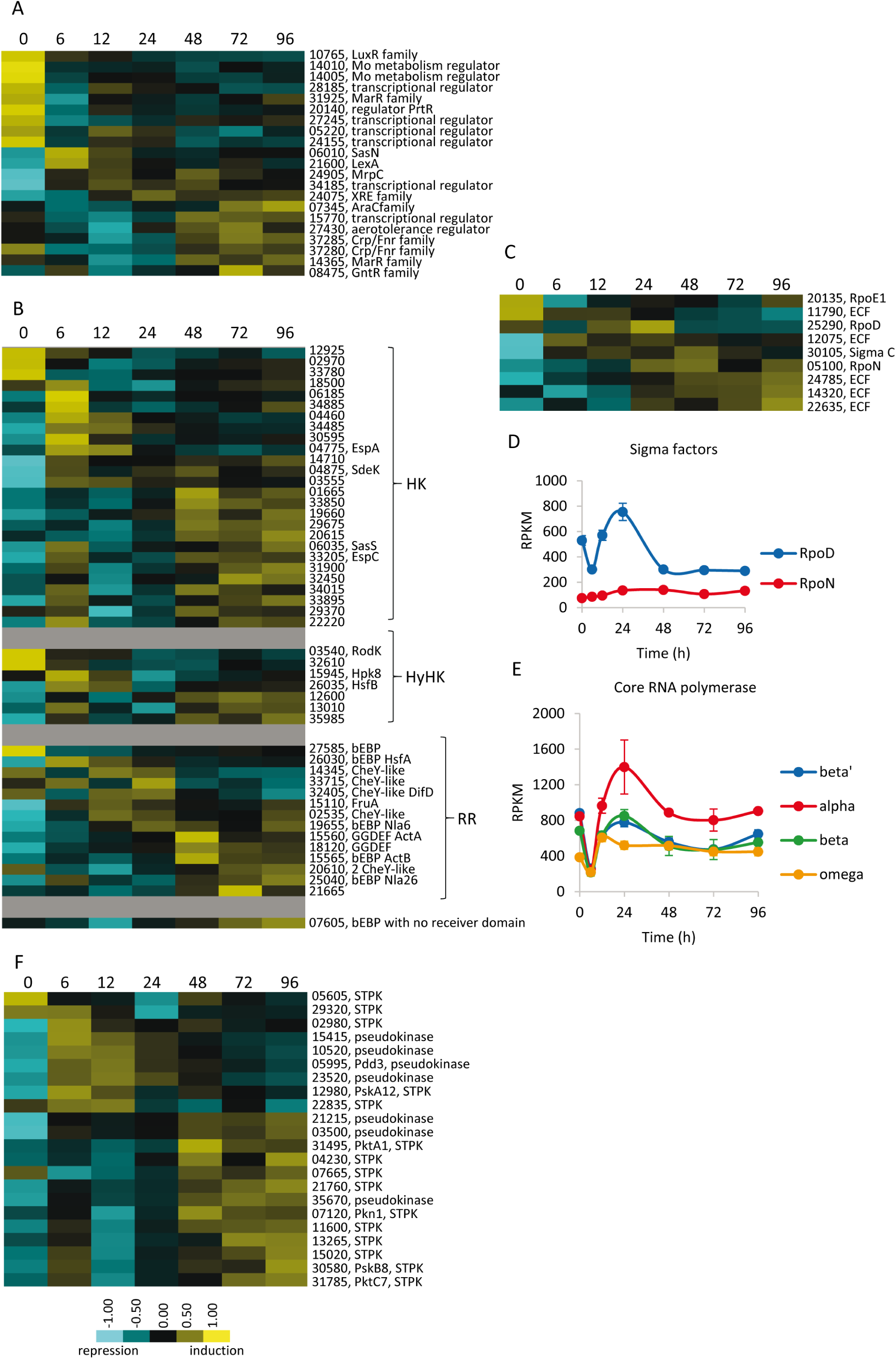
Developmental expression profiles of genes involved in transcriptional regulation and signal transduction. Relative expression levels of genes encoding one-component regulators (A), two component signal transduction proteins (B), sigma factors (C), and serine/threonine protein kinases (F). For simplicity, the MXAN_RS designation was omitted from the locus tag of each gene. Relative expression levels for panels A, B, C, and F are indicated by color code according to the legend at the bottom. Expression levels (RPKM) of genes encoding the major sigma factors (D) and the subunits of the RNA polymerase (E) plotted against developmental time points in hours. Error bars indicate standard deviations.

Of the 272 encoded TCS genes, we found 47 included in the DGs: 26 HKs, 7 hybrid HKs (HyHKs, containing HK and RR modules in the same polypeptide), and 14 RRs. 6 TCS genes were shut down during development (Figure 9B). Only 13 of 47 TCS genes have been previously characterized, thus pinpointing additional candidates for further characterization. Additionally, 5 of the 14 RRs belong to the group of bEBPs (plus MXAN_RS07605, which exhibits an architecture similar to bEBPs, but contains a GAF instead of a receiver domain). bEBPs function to activate expression from sigma 54 dependent promoters (Morett and Segovia, 1993). Besides the bEBPs, FruA is the only other RR found in the DGs that contains a DNA-binding domain (Figure 9B). Out of the remaining 8 RRs, 5 were stand-alone receiver domains (CheY-like), 2 contain a putative diguanylate cyclase output domain, and 1 contained an output domain of unknown function (Figure 9B). As many of the HKs and RRs that were developmentally regulated are orphans, the results presented here may help to identify cognate pairs. Overall, these observations are in good agreement with an established model in which starvation activates four genetic regulatory networks (Kroos, 2016): the bEBP, the Mrp, FruA, and Nla24 modules, with the latter being activated by cyclic-di-GMP (Skotnicka et al., 2016).

Regarding other transcription factors, 9 sigma factors were found to be developmentally regulated in a sequential fashion (Figure 9C), including the major sigma factor RpoD (Inouye, 1990) and RpoN, which encodes sigma 54 (Keseler and Kaiser, 1997). The expression profiles of these two sigma factors clearly differ during development. While *rpoD* was down-regulated at 6 h, up-regulated during aggregation, and then down-regulated during sporulation, *rpoN* is up-regulated throughout development (Figure 9D). The expression profiles of sigma 54 and bEBPs are consistent with previous results demonstrating that a bEBP cascade is triggered upon starvation (Giglio et al., 2011). Additionally, *sigC*, which encodes a group II sigma factor (Apelian and Inouye, 1993), was identified in DG7. The remaining six sigma factors are predicted extracytoplasmic functions (ECF) sigma factors, including RpoE1, involved in motility (Ward et al., 1998). Moreover, RpoE1 and two additional developmentally regulated sigma factors, require the architectural proteins CarD and CarG to activate their own transcription (Abellón-Ruiz et al., 2014). It is noteworthy that the four subunits of the core RNA polymerase are developmentally regulated with profiles similar to that observed for the ribosomal proteins (Figure 9E).

*M. xanthus* encodes 99 STPK, 11 of which have been reported to be pseudokinases, because they lack at least one of the three residues required for phosphorylation (Pérez et al., 2008). 22 STPKs were included in the DGs (Figure 9F). Interestingly, 7 are pseudokinases while 15 are predicted active kinases (Tables S4 and S5), representing 64% and 17% of the total encoded pseudo-and functional kinases, respectively. Considering that most *M. xanthus* active STPKs are RD kinases (Pérez et al., 2008), their activity is predicted to be post-translationally regulated by autophosphorylation (Johnson and Lewis, 2001; Kornev et al., 2006). In contrast, pseudokinases are expected to be unable to autophosphorylate, so they may require regulation via transcription for functional control. Little is known about the developmental role of the STPKs included in the DGs (Muñoz-Dorado et al., 1991; Inouye et al., 2008).

Together, these observations demonstrate that a highly integrated genetic regulatory network directs the developmental program, with some regulators acting simultaneously and others sequentially to perfectly modulate the different events that occur through development.

### Genes identified through chemical induction of sporulation mainly map to DG7 and DG8

*M. xanthus* spore differentiation can be induced in cells growing in rich broth culture by addition of chemicals such as 0.5 M glycerol (Dworkin and Gibson, 1964). This artificial induction process bypasses the requirement for starvation, motility (aggregation), and alternate cell fates (Higgs et al., 2014). Using this process, a core sporulation transcriptome was defined using microarray methodology (Müller et al., 2010). We therefore compared the sporulation transcriptome to this developmental transcriptome and determined 1388 up-, down-, or un-regulated genes described here were included in both transcriptome studies (Figure 2B, Figure 2B—Source data 1 and 2, Figure 2B—figure supplement 1, and Figure 2B—Source data 3). 92% (218/237) of genes significantly up-regulated (>2-fold) in the sporulation transcriptome were also significantly up-regulated in the developmental transcriptome. Up-regulated genes were primarily over-represented in DGs 7 and 8, with low probability (p) that this association is due to random chance (p = 1 x10^−12^ and p = 2 x10^−11^, respectively). Co-regulated genes in these groups include those predicted to be involved in generation of sugar precursors necessary for spore coat and storage compound synthesis (Figure 6). Likewise, genetic loci involved in spore coat synthesis (*exo*) and surface polysaccharide arrangement (*nfs*) (Müller et al., 2012; Holkenbrink et al., 2014) fell in DG8.

As expected, of the large pool of genes (754) that were not significantly regulated in the sporulation transcriptome, a relatively small number (89; 12%) were also not significantly up-or down-regulated in the developmental transcriptome. This pool of genes likely represents constitutively expressed genes that serve as good normalization markers (Figure 2B—Source data 1) and includes housekeeping genes such as the transcription termination factor Rho (MXAN_RS11995) and the cell cytoskeletal protein MreB (MXAN_RS32880) (Müller et al., 2012; Treuner-Lange et al., 2015; Fu et al., 2018), as well as the gliding motility and sporulation protein AglU (MXAN_RS14565) (White and Hartzell, 2000), and the CheA homolog DifE (MXAN_RS32400), which is required for exopolysaccharide production and social motility (Yang et al., 2000). Interestingly, a large number of genes (141) which are not significantly regulated in the sporulation transcriptome, were found enriched in DG10 (p = 7.46 x10^−10^), the latest DG. We speculate this pool of genes may be present in PRs, a cell fate enriched late in the developmental program and not present during chemical induction of sporulation. Proteins encoded by genes in this pool include several STPKs, the bEBP Nla26, and several PKSs. Since PKSs and hybrid NRPS/PKS are up-regulated during the latest stages of starvation development but are down-regulated or non-regulated in chemical-induced spores (Tables S4, S5, and S6), it is tempting to speculate that the SMs are induced in the PRs to defend the spores inside the fruiting bodies.

### Toward elucidation of a complete developmental gene-regulatory circuit

The transcriptomic analyses presented here both corroborate the data obtained by myxobacteriologists on *M. xanthus* in the last 40 years and offer some new explanations about how this myxobacterium undergoes its developmental program. Although this developmental cycle was considered one of the most complex among prokaryotes, the high number of genes found in this study to be developmentally regulated at the mRNA level indicates that this complexity is much higher than expected. Undoubtedly, the developmental mRNA expression profiling presented here will act as a blueprint for the complete elucidation of the *M. xanthus* developmental regulatory program.

## MATERIALS AND METHODS

### Preparation of cells for RNA-Seq experiment

*M. xanthus* strain DK1622 (Kaiser, 1979; Goldman et al., 2006) was used in this study. Cells were grown in CTT liquid medium (Hodgkin and Kaiser, 1977) at 30°C with vigorous shaking (300 rpm) to 3.0×10^8^ cells/ml (optical density at 600 nm [OD_600_] of 1), and then harvested and resuspended in TM buffer (10 mM Tris-HCl [pH 7.6], 1 mM MgSO_4_) to a calculated density of 4.5×10^9^ cells/ml (OD_600_ of 15). For each time replicate, 200 μl aliquots of concentrated cell suspension were spotted onto thirteen separate CF agar plates (Hagen et al., 1978). Two replicates of cells were harvested from plates at 6, 12, 24, 48, 72 and 96 h (samples WT_6, WT_12, WT_24, WT_48, WT_72 and WT_96, respectively), and the obtained pellets were transferred immediately into 0.5 ml of RNA Protect Bacteria Reagent (Qiagen). Cells were then incubated at room temperature for 5 min, harvested by centrifugation at 5000 g for 10 min (4°C), and stored at - 80°C after removal of the supernatant. For the t=0 samples (sample WT_0), two replicates of 30 ml of the original liquid culture (OD_600_ of 1) were harvested by centrifugation as above, resuspended in RNA Protect Bacteria Reagent, and processed in the same manner.

### RNA extraction

To isolate RNA, frozen pellets were thawed and resuspended in 1 ml of 3 mg/ml lysozyme (Roche) and 0.4 mg/ml proteinase K (Ambion) prepared in TE buffer (10 mM Tris-HCl; 1 mM ethylenediaminetetraacetic acid [EDTA], pH 8.0) for cell lysis. Samples were incubated 10 min at room temperature. RNA extraction was carried out using the RNeasy Midi Kit (Qiagen) and each sample was eluted in 300 µl of RNase-free water. The concentration of RNA was measured using a NanoDrop ND-2000 spectrophotometer (NanoDrop Technologies, USA). To remove DNA, each RNA sample was supplemented with 1 unit of DNase I (from the DNA amplification grade Kit of Sigma) per µg of RNA and incubated at room temperature for 10 min. The reaction was stopped by adding the stop solution included in the kit and incubating 10 min at 70°C. The obtained RNA was precipitated with 1/10 volume of 3 M sodium acetate and 3 volumes of ethanol, and resuspended in 50 µl of RNase-free water. The quality of the total RNA was verified by agarose gel electrophoresis, and the concentration was determined using NanoDrop as indicated above.

### Double stranded copy DNA synthesis

First strand DNA was synthesized using SuperScript III Reverse transcriptase (Invitrogen) starting with 5 μg of RNA in a final reaction volume of 20 µl. In the next step, second-strand DNA was synthesized by adding 40 units of *E. coli* DNA polymerase (New England Biolabs), 5 units of *E. coli* RNAse H (Invitrogen), 10 units of *E. coli* DNA ligase (New England Biolabs), 0.05 mM (final concentration) of dNTP mix, 10x second-strand buffer (Biolab), and water to 150 µl. After 2 h at 16°C, the reaction was stopped with 0.03 mM EDTA (final concentration). The obtained DNA was purified and concentrated using the DNA Clean and Concentrator Kit of Zymo Research according to manufacturer’s instructions. The final product was eluted in DNA elution buffer from the kit to reach, at least, a yield of 2 µg of DNA, with a minimum concentration of 200 ng/µl.

### Sequencing and transcriptomic data analysis

The cDNA from two biological replicates of each condition (see above) was used for sequencing using the Illumina HiSeq2000 (100-bp paired-end read) sequencing platform (GATC Biotech, Germany). Sequence reads were pre-processed to remove low-quality bases. Next, reads were mapped against *M. xanthus* DK1622 ribosomal operon sequences using BWA software with the default parameters (Li and Durbin, 2009). Remaining reads were subsequently mapped to the genome sequence with the default parameters and using the pair-end strategy. SAMtools (Li et al., 2009) were used to convert resulting data into BAM format. Artemis v.16.0.0 (Rutherford et al., 2000) was used for the visualization of the sequence reads against the *M. xanthus* genome. Once the transcripts were mapped to the genome, the average median value for each condition was used in further analyses (Tables S1 and S2).

### Developmental gene analysis

Genes with fewer than 50 reads in a given time point were removed from analysis. RPKM values of the remaining genes were then compared across the developmental time-points. Developmental expression was characterized by a >2-fold change in RPKM values across the time course and a >0.7 R^2^ correlation coefficient of the two time course replicates. Genes passing these criteria are present in Tables S3 and S4. Randomization of the time point RPKM values within each replicate data set yielded a false discovery rate of 3.97% based on 5 randomized simulations that scrambled the order of the time points across all genes in the two datasets. Genes passing the developmental expression criteria were hierarchically clustered using the kmeans algorithm with 10 DGs using the cluster 3.0 software package (Hoon et al., 2004).

### Assay of β-galactosidase activity

For quantitative determination of β-galactosidase activity during development, strains containing *lacZ* fusions (Supplementary file 1) were cultured and spotted onto CF plates as described above. Cell extracts were obtained at different times by sonication and assayed for activity as previously reported (Moraleda-Muñoz et al., 2003). The amount of protein in the supernatants was determined by using the Bio-Rad protein assay (Bio-Rad, Inc.) with bovine serum albumin as a standard. Specific activity is expressed as nmol of *o*-nitrophenol produced per min and mg of protein. The results are the average and associated standard deviation from three independent biological replicates.

### Supporting data

The transcriptome sequencing data (raw-reads) was submitted to NCBI SRA under the Bioproject accession number: PRJNA493545. SRA accession numbers for each of the replicas are as follows: 0 h: SAMN10135973 (WT_0_1-biological_replicate_1) and SAMN10135974 (WT_0_2-biological_replicate_2); 6 h: SAMN10135975 (WT_6_1-biological_replicate_1) and SAMN10135976 (WT_6_2-biological_replicate_2); 12 h: SAMN10135977 (WT_12_1-biological_replicate_1) and SAMN10135978 (WT_12_2-biological_replicate_2); 24 h: SAMN10135979 (WT_24_1-biological_replicate_1) and SAMN10135980 (WT_24_2-biological_replicate_2); 48 h: SAMN10135981 (WT_48_1-biological_replicate_1) and SAMN10135982 (WT_48_2-biological_replicate_2); 72 h: SAMN10135983 (WT_72_1-biological_replicate_1) and SAMN10135984 (WT_72_2-biological_replicate_2); 96 h: SAMN10135985 (WT_96_1-biological_replicate_1) and SAMN10135986 (WT_96_2-biological_replicate_2).

## ACKNOWLEDGEMENTS

This work has been supported by the Spanish Government (grants CSD2009-00006 to José Muñoz-Dorado and BFU2016-75425-P to Aurelio Moraleda-Muñoz (70% funded by FEDER), and by NIGMS of the National Institutes of Health under award number R35GM124733 to JMS. JPT and JMD were granted with fellowships of the Salvador de Madariaga Program to stay at WSU for four months. PIH was funded by a grant from NSF IOS 1651921. We also want to thank Prof. Lee Kroos for providing strains.

## SUPPLEMENTS

**Table 1—Source data 1.** Number of reads for each ORF of *Myxococcus xanthus* at 0, 6, 12, 24, 48, 72 and 96 hours of development.

**Table 1—Source data 2.** RPKM values of the developmental time course and correlation scores. RPKM values reported here were calculated from the total number of non-tRNA/rRNA containing reads. The old (MXAN_) locus tags, new gene identifiers (MXAN_RS), gene name and predicted functions or pathways in which they have been previously implicated are included. The number of missing data points and fold change were used as criteria for the developmental gene analysis. DG or reason that genes were not included in the DGs is indicated.

Supplementary file 1. Validation of the RNA Seq transcription patterns for genes *spiA* (MXAN_RS2076) and *fmgE* (MXAN_RS16790) (B). β-galactosidase specific activity (SA) of the strains harboring *lacZ* fusions to the respective genes (red lines) compared to RNA seq RPKM values (blue lines) at each developmental time point (h). Error bars indicate standard deviations.

**Figure 2A—Source data 1.** RPKM values for developmentally controlled genes and distribution of the genes in the ten development groups. Genes with >2-fold change in mRNA levels and >0.7 correlation between the two replicate developmental time courses are listed with their corresponding RPKM mRNA level measurements, following the same order shown in Figure 2. The old (MXAN_) locus tags, new gene identifiers (MXAN_RS), gene name and predicted functions or pathways in which they have been previously implicated are included.

**Figure 2A—Source data 2.** Previously reported developmental genes and identification in the ten developmental groups. Genes have been highlighted by color (see color code) to depict known or implied roles in developmental processes and/or membership of protein families involved in development. References are the same as in Figure 2A—Source data 1.

**Figure 2B—Source data 1.** Genes included for comparison of developmental and sporulation transcriptomes. Developmental transcriptome genes (this study) include those assigned to DGs and those which passed requirements for all data points with >0.7 correlation (considered constitutive). Sporulation transcriptome data were from Müller et. al. (2010) BMC Genomics 11:264, in which vegetative cells were induced to artificially sporulate by addition of glycerol to M. The sporulation gene data set included genes significantly down-, up-, or not regulated. Up-regulated genes were classified as up1 (peak expression within 2 h of sporulation induction) or up2 (peak expression at 2-4 h of sporulation induction). Only genes present in both studies are included in this table.

**Figure 2B—Source data 2.** Tally of enrichment of sporulation transcriptome genes [Müller et. al. (2010) BMC Genomics 11:264] found in each of the DGs. Only genes identified as having reliable expression patterns in both developmental and sporulation transcriptome studies (Figure 2B— Source data 1) were included. Spore transcriptome genes enriched or depleted in any DG were tested for significance (p < 0.01) using a chi square test [as per Müller et. al. (2010) BMC Genomics 11:264] generating probability (p) values that this enrichment was due to chance. Sporulation transcriptome up-, down-or not regulated gene categories significantly over-enriched in any DGs are highlighted in yellow, genes significantly depleted from any DG category are highlighted in blue. Text in red: sporulation up-regulated genes are enriched in DGs 7 and 8.

**Figure 2B—Source data 3.** Tally of genes observed in both the developmental (this study) and sporulation [Müller et. al. (2010) BMC Genomics 11:264] transcriptomes as supporting data for Figure 2B—figure supplement 1. Total number indicates genes with expression profiles considered reliable in both data sets. Developmental DG1 was considered down-regulated genes, while up-regulated genes were considered all genes in DGs2-10. Developmental constitutive genes were those whose expression level was not more than 2-fold different from vegetative cells at any time point.

**Figure 2B – figure supplement 1.**
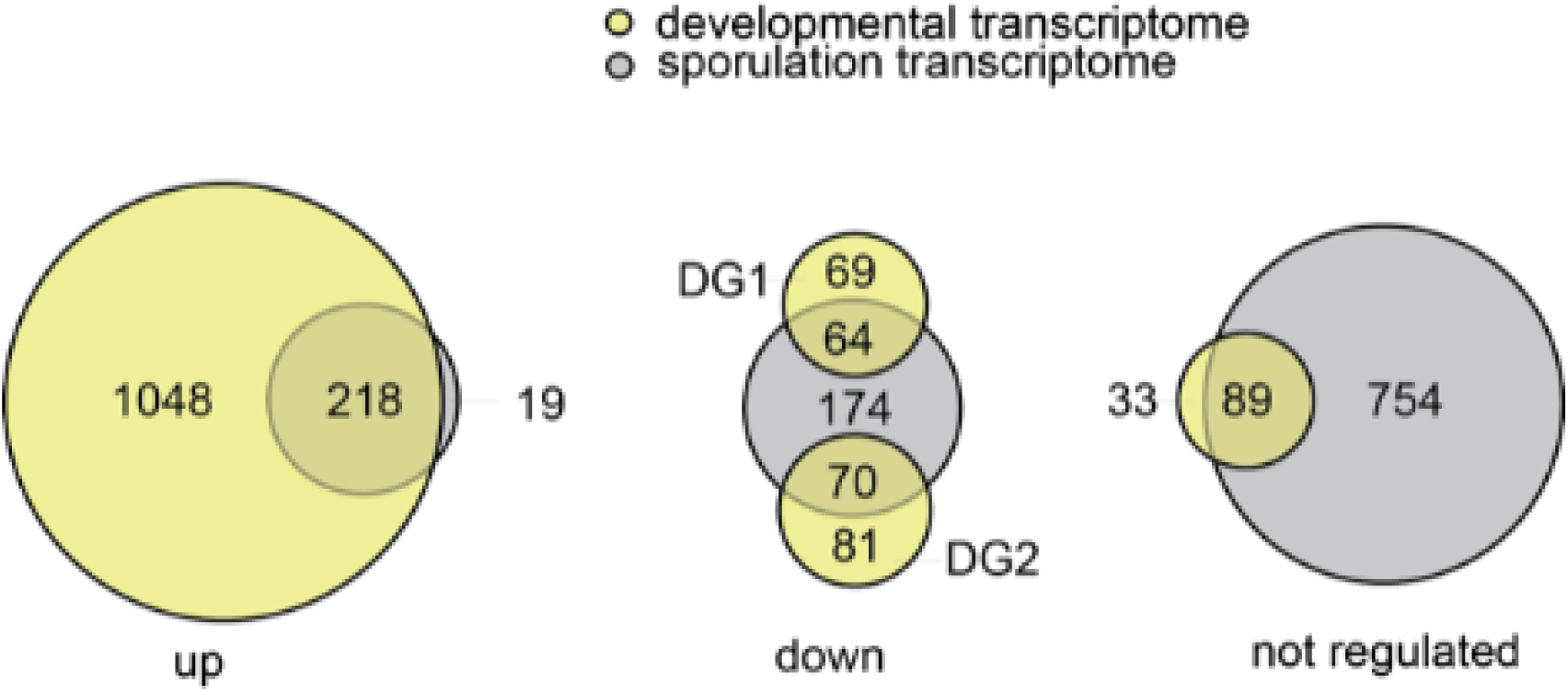
Distribution of co-regulated genes observed in sporulation [Müller et. al., (2010) BMC Genomics 11:264] and developmental transcriptomes (this study). Venn diagrams illustrating genes considered significantly up-(left), down (middle), or not regulated (left) in the developmental (yellow) and sporulation (grey) transcriptome studies. Genes showing the same expression patterns are represented by overlapping circles, while genes with expression patterns specific to either process are indicated in non-overlapping. Circle size is proportional to the number of genes observed in each category. Only genes with expression profiles that passed correlation criteria available in both studies (Figure 2B—Source data 1) were included. Genes considered developmentally down-regulated were from DG1 and DG2. Supporting tally information in Figure 2B—Source data 2.

**Supplementary Fig 1.**
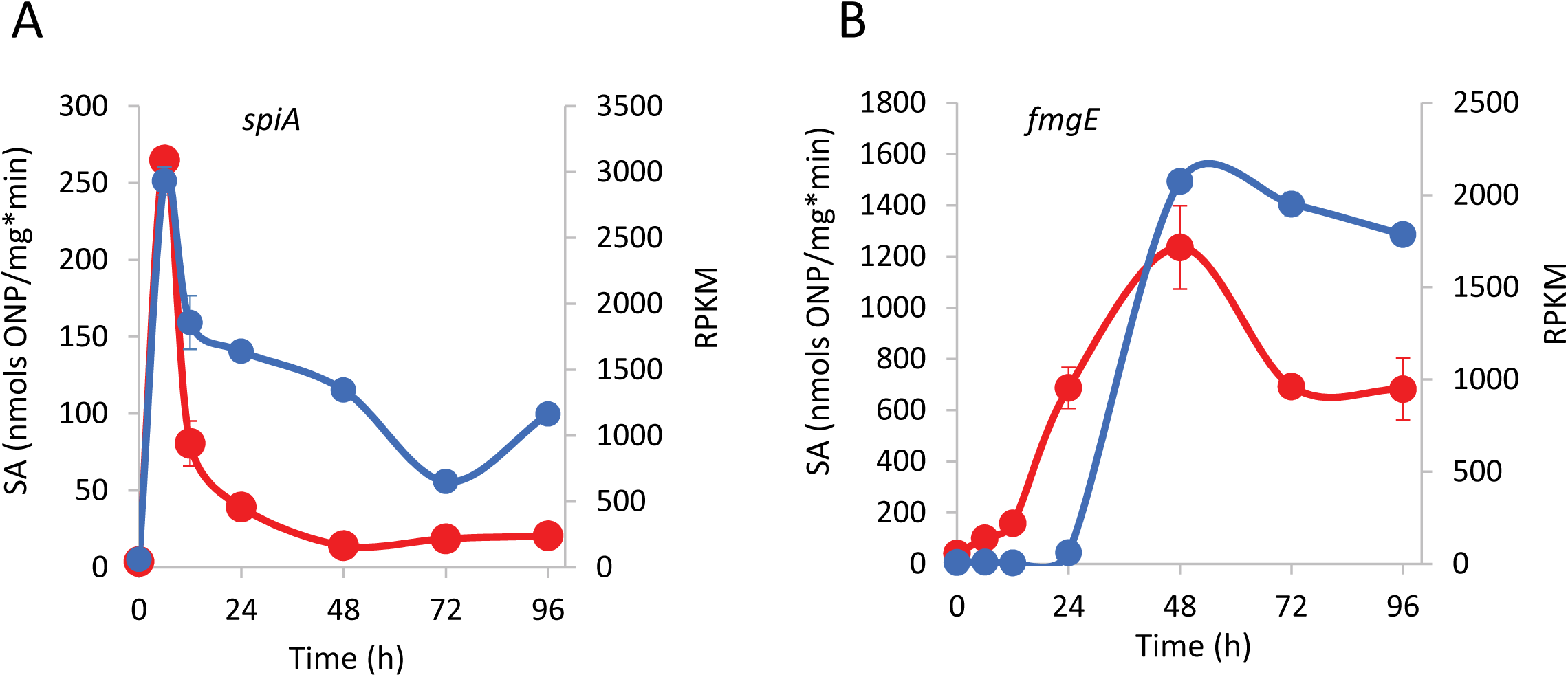
Validation of the RNA Seq transcription patterns for genes spiA (MXAN_RS2076) (A) and fmgE (MXAN_RS16790) (B). β-galactosidase specific activity (SA) of the strains harboring lacZ fusions to the respective genes (red lines) compared to RNA seq RPKM values (blue lines) at each developmental time point (h). Error bars indicate standard deviations.

Bacterial strains used to validate RNA Seq expression profiles using β-galactosidase activity produced by *lacZ* fusions.

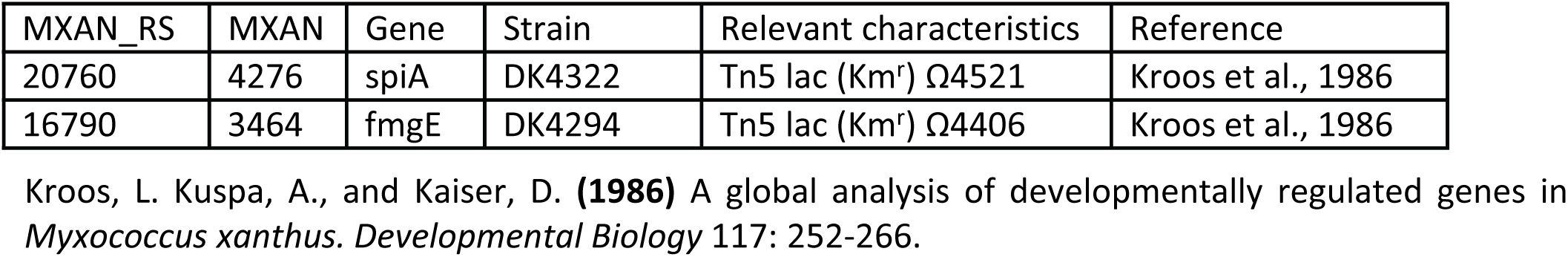

